# Computational complexity as an ultimate constraint on evolution

**DOI:** 10.1101/187682

**Authors:** Artem Kaznatcheev

## Abstract

Experiments show that evolutionary fitness landscapes can have a rich combinatorial structure due to epistasis. For some landscapes, this structure can produce a computational constraint that prevents evolution from finding local fitness optima – thus overturning the traditional assumption that local fitness peaks can always be reached quickly if no other evolutionary forces challenge natural selection. Here, I introduce a distinction between easy landscapes of traditional theory where local fitness peaks can be found in a moderate number of steps and hard landscapes where finding local optima requires an infeasible amount of time. Hard examples exist even among landscapes with no reciprocal sign epistasis; on these semi-smooth fitness landscapes, strong selection weak mutation dynamics cannot find the unique peak in polynomial time. More generally, on hard rugged fitness landscapes that include reciprocal sign epistasis, no evolutionary dynamics – even ones that do not follow adaptive paths – can find a local fitness optimum quickly. Moreover, on hard landscapes, the fitness advantage of nearby mutants cannot drop off exponentially fast but has to follow a power-law that long term evolution experiments have associated with unbounded growth in fitness. Thus, the constraint of computational complexity enables open-ended evolution on finite landscapes. Knowing this constraint allows us to use the tools of theoretical computer science and combinatorial optimization to characterize the fitness landscapes that we expect to see in nature. I present candidates for hard landscapes at scales from single genes, to microbes, to complex organisms with costly learning (Baldwin effect) or maintained cooperation (Hankshaw effect). Just how ubiquitous hard landscapes (and the corresponding ultimate constraint on evolution) are in nature becomes an open empirical question.

---

Genotype and fitness are two central concepts in evolutionary biology. Through its production of phenotypes and those phenotypes’ interactions with the biotic and abiotic environment, a given genotype has a certain fitness. A fitness landscape summarizes this relationship between genotypes (or phenotypes) and fitness. Formally, fitness landscapes combine numeric fitnesses and a mutation-graph into a combinatorially structured space where each vertex is a possible genotype (or phenotype). The numeric structure is given by a function that maps each genotype to a fitness; typically represented as a non-negative real number and having different physical operationalizations in different experimental systems. A given genotype is more similar to some rather than other genotypes – giving us a notion of genetic distance or mutation-graph. The mutation-graph specifies which genotypes are similar, typically as edges between any two genotypes that differ in a single mutation. This provides the combinatorial structure. A genotype is a local fitness peak (or local fitness optimum) if no adjacent genotype in the mutation-graph has higher fitness.

We usually imagine fitness landscapes as hills or mountain ranges, and continue to assume – as Wright [1] originally did – that on an arbitrary landscape “selection will easily carry the species to the nearest peak”. And we define a *constraint* as anything that keeps a population from reaching a local fitness peak. For those that view evolution as a sum of forces, with natural selection being only one of them, it is possible for other forces to act as a constraint when they overpower natural selection and keep the population away from a local fitness peak. Such cases are often associated with maladaptation [2] and are usually attributed to mechanisms like mutation meltdown, mutation bias, recombination, genetic constraints due to lack of variation, or explicit physical or developmental constraints of a particular physiology. I will refer to such situations, where non-selection forces (and/or aspects internal to the population) keep the population from reaching a local fitness peak, as *proximal* constraints on evolution. In contrast, I will refer to a constraint as *ultimate* if it is due exclusively to features of the fitness landscape and is present in the absence of other forces or even holds regardless of the strength of other forces. All constraints are either proximal, ultimate, or a mix of the two. I introduce this terminology of proximal and ultimate *constraints* by analogy to Mayr’s distinction between proximal and ultimate *causes* in biology [3]. Mayr considered as ultimate only those evolutionary causes that are due exclusively to the historic process of natural selection [4], so I consider as ultimate only those evolutionary constraints that are due exclusively to the fitness landscape structure of natural selection.

The distinction that I am making between proximal and ultimate constraints can be made clearer by reference to a distinction in computer science between algorithms and problems. I will consider the population structure, update rules, developmental processes, mutation operator or bias, etc as together specifying the algorithm that is evolution. In contrast, the families of fitness landscapes are like problems to be ‘solved’ by evolution and specific fitness landscapes on which populations evolve are problem-instances. A *proximal constraint* is any feature of the evolutionary algorithm that prevents the population from finding a local fitness optimum in polynomial time. For a classic example, consider a population with an extreme lack of genetic variation that cannot proceed to an adjacent fitter genotype because the allele that it differs in is simply not available in the population. In this case, the proximal constraint of lack of variation due to the details of this particular population’s evolutionary algorithm prevents it from reaching a fitness peak. In contrast, an *ultimate constraint* is any feature of the problem (i.e. family of fitness landscapes) that prevents the population from finding a local fitness optimum in polynomial time. It is the goal of this paper to show a convincing example of such a constraint.

One candidate for an ultimate constraint on evolution – historicity or path-dependence – is already widely recognized. A local peak might not be the tallest in the mountain range, so reaching it can prevent us from walking uphill to the tallest peak. This constraint has directed much of the work on fitness landscapes toward how to avoid sub-optimal peaks or how a population might move from one peak to another [1,5]. Usually, these two questions are answered with appeals to the strength of other evolutionary forces. Although sometimes the second question is sidestepped by postulating that local fitness peaks are part of the same fitness plateau in a holey adaptive landscape and thus fitness valleys can be bypassed to move between different local optima in the plateau [6,7]. But both of these types of questions implicitly assume that local peaks (or plateaus) are easy to reach and thus the norm for natural selection. When the constraint of historicity is active, being at one local optimum prevents the population from reaching other (higher) local optima. Thus, this candidate for an ultimate constraint is only partial: it prevents only certain – not all – local fitness optima from being found. In this case, it prevents evolution from finding the highest local peak: the global optimum. But, we seldom consider that even reaching *any* local optimum might be impossible in a reasonable amount of time.

Here I show that computational complexity is an ultimate constraint on evolution: it can prevent evolution from finding *any* local fitness peak – even low fitness ones. A careful analysis – formal mathematical proofs for all statements are available in the supplementary appendix (SA; for a summary, see SA Table 1) and referenced throughout the text – shows that the combinatorial structure of fitness landscapes can prevent populations from reaching any local fitness peaks. This suggests an alternative metaphor for fitness landscapes: fitness landscapes as mazes with the local fitness optima as exits. Natural selection cannot see far in the maze and must rely only on local information from the limited genetic variation of nearby mutants. In hard mazes, we can end up following exponentially long winding paths to the exit because we cannot spot the shortcuts. In such cases, even if natural selection is the only force acting on the population, a fitness optimum cannot be found. Worse yet, the hardest mazes might not have any shortcuts and even the most clever and farsighted navigator will not know how to reach an exit in a feasible amount of time. In other words, even if the other evolutionary forces ‘conspire to help’ natural selection, a local fitness optimum cannot be found.

**Table 1:**
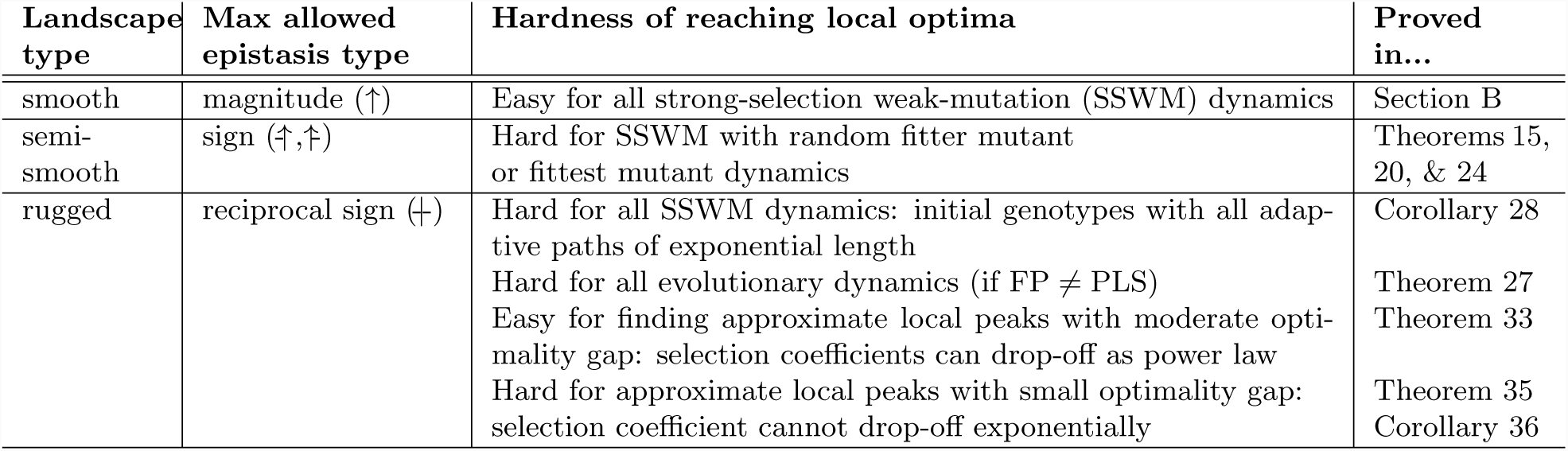
Summary of main results. Each landscape type (column 1) is characterized by the most complicated permitted type of epistasis (column 2; see A.1). Based on this, there are families of this landscape type that are easy or hard under progressively more general dynamics (column 3), which is proved in the corresponding part of the appendix (column 4).

To establish these results, I will introduce into biology new techniques from theoretical computer science for managing the complexity of fitness landscapes. The best current techniques in biology come from the statistical mechanics of disordered systems and rely on “[t]he idea that unmanageable complexity can be replaced by randomness” [8]. This statistical approach uses randomness in two places: (1) the random mutations, birth-death events, and other physical and biological processes within the algorithm of evolution, and (2) the theoretical distributions of fitness landscapes themselves. For a computer scientist, the first use of randomness corresponds to the analysis of randomized algorithms – certainly a good decision when thinking about evolution. The second use of randomness corresponds to average-case analysis over problem instances. But when the real-world distribution of problem-instances in unknown or hard to characterize, computer scientists are hesitant to pick a specific simple distribution just to analyze the algorithm. Instead, computer scientists usually specify a formal, logically-defined hypothesis class of conceivable problem-instances and then analyze their algorithm for arbitrary distributions over these instances.

In this report, I embrace the randomness within the algorithm – i.e. the randomness of evolution. But instead of introducing a convenient-to-analyze distribution of possible fitness landscapes, I focus on worst-case analysis. In this way, this report can be seen as a contribution to the small but growing literature on population genetics and evolutionary biology through the algorithmic lens [9–16].

By focusing on worst-case analysis, I am constructing – sometimes implicitly – families of fitness landscapes that are consistent with the logical structure of our hypothesis class of conceivable fitness landscapes. I then show that in these hard fitness landscapes, computational complexity is an ultimate constraint. But this should not be interpreted as a claim that hard landscapes are ubiquitous or that computational complexity is a *major* constraint. That would be an empirical question that depends on which fitness landscapes occur in nature. In this report, I suggest several candidates that I suspect correspond to hard landscapes, but the general empirical question of ubiquity is beyond the scope of this theoretical work.

## Epistasis & semi-smooth landscapes

What makes some fitness landscapes difficult to navigate is that the effects of mutations at different loci interact with each other. Epistasis is a measure of the kind and amount of inter-locus interactions. If the fitness effect of a mutation *a → A* can have a different sign depending on the genetic background *b* or *B* of another locus then these two loci are said to have sign epistasis (Figure 1b and SA Definition 5). If both mutations have one sign on their own, but the opposite sign together – either bad + bad = good or good + good = bad – then the landscape has reciprocal sign epistasis ([17– 19]; Figure 1c and SA Definiton 6). A classic example of reciprocal sign epistasis is a lock-and-key, changing just one of the lock or the key breaks the mechanism, but changing both can be beneficial. Finally, magnitude epistasis (positive and negative; SA Definition 4) are inter-locus interactions that deviate from additivity, but do not change the sign of fitness effects. This type of epistasis does not change the combinatorial structure of the landscape. As such, I treat it simply as a lack of sign-epistasis.

**Figure 1:**
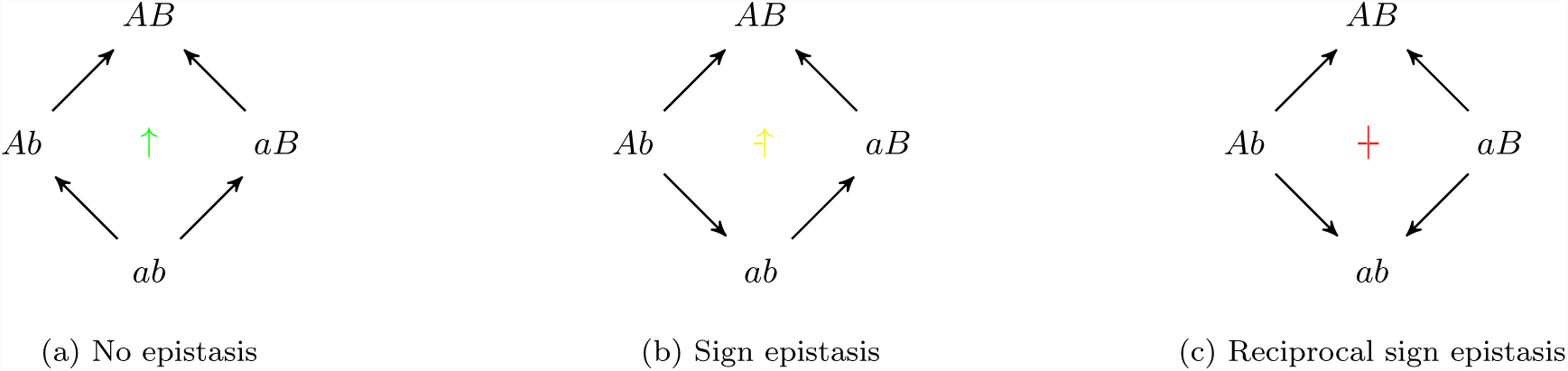
Three different kinds of epistasis possible in fitness graphs. Arrows are directed from lower fitness genotypes towards mutationally adjacent higher fitness genotypes. Genes *a, A* and *b, B* are labeled such that fitness *w*(*AB*) *> w*(*ab*). In the center of each graph is a marker for the type of epistasis, the marker’s various rotations & reflections cover the cases where *AB* does not have the highest fitness. For this more exhaustive classification and discussion see SA Figure 4 & SA A.1

A landscape without sign epistasis – like the *Escherichia coli β*-lactamase fitness landscape measured by Chou, Chiu, Delaney, Segre, and Marx [20] in figure 2a – is called *smooth* ([19, 21] and SA B), so let’s call a fitness landscape *semismooth* if it has no reciprocal sign epistasis. The fitness graphs ([19] and SA A) of semi-smooth fitness landscapes are equivalent to acyclic unique sink orientations previously defined in a different context by Szabó and Welzl [22] for the analysis of simplex algorithms (SA Definition 12 and Proposition 13). Since reciprocal sign epistasis is a necessary condition for multiple peaks (SA Corollary 11 and Poelwijk, Sorin, Kiviet, and Tans [18]), both smooth and semi-smooth fitness landscapes have a single peak *x*^∗^. Further, there are short adaptive paths in both: from any genotype *x* there always exists some adaptive path to *x*^∗^ of length equal to the number of loci on which *x* and *x*^∗^ differ (SA Theorem 14). This means that an omniscient navigator that always picks the ‘right’ adaptive point-mutation can be guaranteed to find a short adaptive path to the peak. But unlike smooth landscapes, in a semi-smooth landscape not every shortest path is adaptive and not every adaptive path is short. And since evolution does not have the foresight of an omniscient navigator, it is important to check which adaptive path myopic evolutionary dynamics will follow.

**Figure 2:**
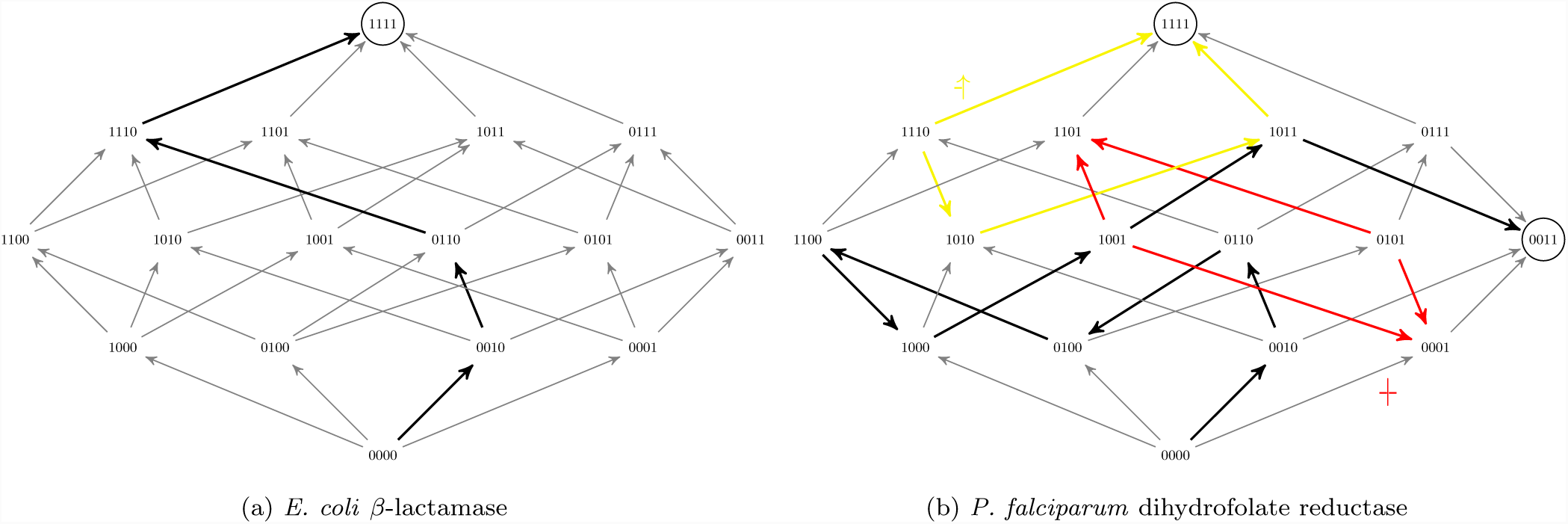
Two examples of empirical biallelic fitness landscapes on four loci. Arrows are directed from lower fitness genotypes to higher and fitness optima are circled. Examples of adaptive dynamics are highlighted with thick black arrows. Figure 2a, based on the *E. coli β*-lactamase data of Chou, Chiu, Delaney, Segre, and Marx [20], is a smooth landscape with no sign epistasis. Thus, it contains a single optimum (1111). Figure 2b is based on Lozovsky, Chookajorn, Brown, et al. [23]’a *P. falciparum* dihydrofolate reductase growth rate data in the absence of pyrimethamine. It has two peaks (0011 & 1111) and both single sign (an example in yellow; -↑) and reciprocal sign epistasis (example in red; †). Based on Szendro, Schenk, Franke, Krug, and de Visser [24]’s Figure 1.

When mutation is weak, we can assume that a population is always monomorphic except for a brief moment of transition as a new mutant fixes. Thus, we can represent the population as a single point on the fitness landscape with an evolutionary step corresponding to a selective sweep that moves the population to a neighbouring genotype. When selection is strong, we can further assume that the evolutionary step takes us to a neighbouring genotype of higher fitness. The rule for selecting which neighbour ends up fixing depends on the details of our mutation operator and model of evolution; i.e. this rule specifies the algorithm. I will refer to the set of algorithms corresponding to all such rules as strong-selection weak mutation (SSWM) dynamics. A number of rules (or algorithms) have been suggested for which fitter neighbour will take over the population [25] – such rules correspond to different models of evolution. The two most common rules are to select a fitter mutant uniformly at random, or to select the fittest mutant. These rules capture the intuition of evolution proceeding solely by natural selection with other forces absent or negligible. All SSWM rules will quickly find the fitness optimum in a smooth fitness landscape. But there exist semi-smooth fitness landscapes such that when starting from a random initial genotype, an exponential number of evolutionary steps will be required for either the random fitter mutant ([26]; SA Theorem 15) or fittest mutant (SA Theorems 20 and 24) dynamics to find the unique fitness optimum. For a small example on six loci, see Figure 3: the black arrows trace the evolutionary path that a population would follow under fittest mutant SSWM dynamics. Although two step adaptive paths exist to the fitness peak (like 000000 *→* 000001 *→* 000011), the myopic navigator cannot notice these shortcuts and ends up on a long winding path. In other words, even when there is a single peak and adaptive paths of minimal length to it, SSWM dynamics can take exponential time to find that peak.

**Figure 3:**
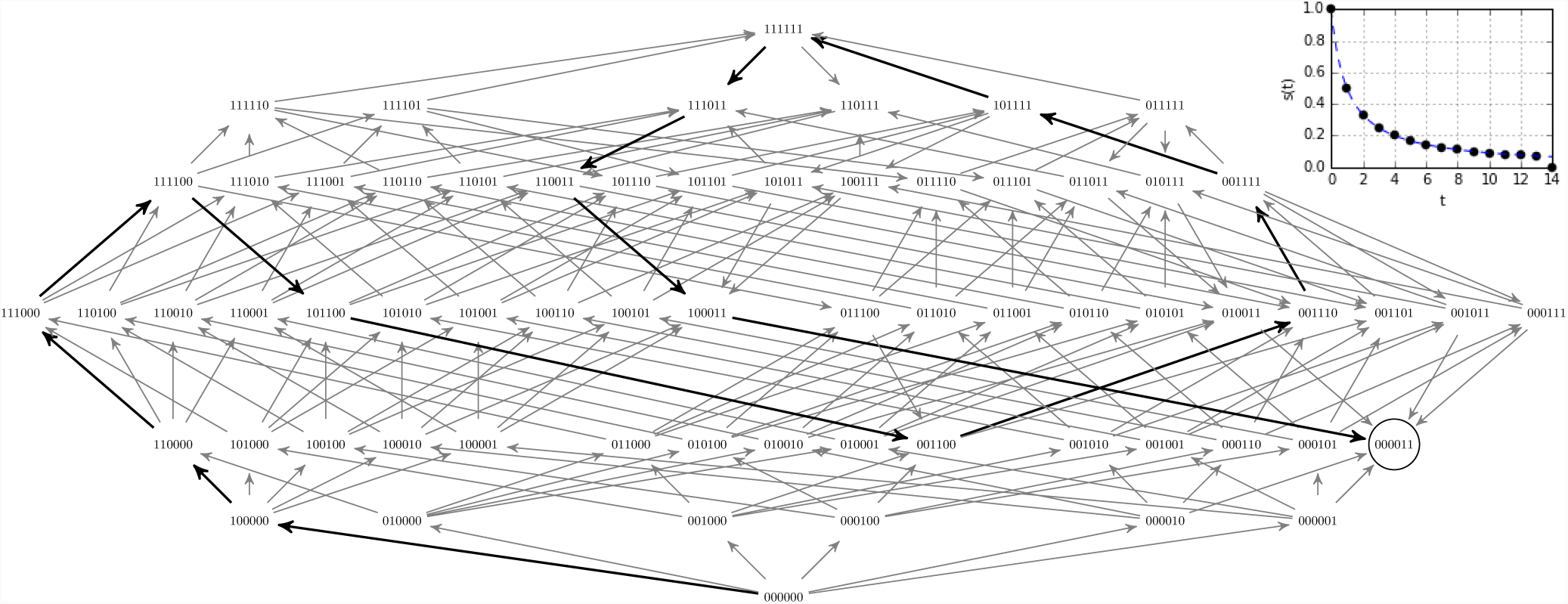
Fittest mutant adaptive path in a winding semi-smooth fitness landscape. An example on six loci of the winding semi-smooth fitness landscapes from SA C.2 on which the length of the path followed by fittest-mutant SSWM dynamics scales exponentially with the number of loci. Here the black arrows are the fittest available mutation, and the adaptive path takes 14 steps to reach the fitness peak at 000011. For the generalization of this landscape to 2*n*-loci, it would take 2^*n*+1^ *-*2 steps for fittest mutant dynamics to reach the fitness peak at (00)^*n-*1^11 (SA Theorem 20). Inset is the selection coefficient 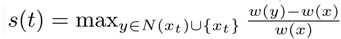; SA D.2) versus mutation step number (*t*) for the fittest mutant adaptive path.

These results show that the computational complexity of the combinatorial structure can be enough to stop evolution from reaching a fitness optimum within a reasonable timescale, even in the absence of suboptimal local peaks. Computer scientists have found it helpful to distinguish between processes that require a time that grows polynomially with the size of the input – generally called tractable – and those that require a time that increases faster than any polynomial (super-polynomial) – intractable. If the winding fitness landscapes of Figure 3 is generalized to 2*n* loci instead of just 6 (SA C.2) then following fittest mutant SSWM dynamics to the peak is an intractable process since it scales exponentially, requiring 2^*n*+1^ −2 mutational steps. Although evolutionary time is long, it is not reasonable to think of it as exponentially long. For example, the above winding process with a genotype on just 120 loci and with new set of point-mutants and selective sweep at a rate of one every second would require more seconds than the time since the Big Bang.

To capture this infeasibility of super-polynomial scaling in time, I introduce a distinction between *easy* and *hard* families of fitness landscapes. If we can guarantee for any landscape in the family that a local fitness peak can be found by natural selection in a time the scales as a polynomial in the number of loci – as is the case for smooth fitness landscapes – then I will call that an *easy* family of landscapes. I will call a family of landscapes *hard* if we can show that the family contains landscapes where finding a local fitness optimum requires a super-polynomial amount of time – as I showed above for semi-smooth fitness landscapes. Given that even for moderately sized genomes such large times are not realizable even on cosmological timescales, I will use “impossible” as a shorthand for “requiring an infeasible amount of time”.

Given their exponential size, it is impossible to completely measure whole fitness landscapes on more than a few nucleotides. But with improvements in high-throughput second-generation DNA sequencing there is hope to measure local fitness landscapes of a few mutations away from a wildtype. Puchta, Cseke, Czaja, Tollervey, Sanguinetti, and Kudla [27] estimated the fitness of 981 single-step mutations of a 333-nucleotide small nucleolar RNA (snoRNA) gene in yeast. They found no neighbours fitter than the wild type gene, this suggests that this gene is already at a fitness peak, and hence that the snoRNA gene’s fitness landscape is easy. In contrast, Li, Qian, Maclean, and Zhang [28] estimated the fitness of 207 single-step mutants of a 72-nucleotide transfer RNA (tRNA) gene, also in yeast, finding two neighbours that are significantly fitter than the wildtype and a number that are fitter but only within experimental noise. Thus, the wildtype tRNA gene is apparently not at a local fitness peak, and suggests this system as a candidate for hard fitness landscapes. Both studies also looked at many 2-and 3-step mutants, and the landscape of the tRNA gene was measured to have more than 160 cases of significant sign epistasis [28], with none in the snoRNA landscape [27], mirroring the difference between hard semi-smooth fitness landscapes and easy smooth landscapes that I am proposing here.

### Rugged landscapes & approximate peaks

But there exist natural fitness landscapes that are even more complicated than semi-smooth ones. For example, we know that some landscapes can contain reciprocal sign epistasis like the Lozovsky, Chookajorn, Brown, et al. [23] *Plasmodium falciparum* dihydrofolate reductase fitness landscape in figure 2b. This is a rugged fitness landscape with two distinct fitness peaks at 0011 and 1111. Although there is not enough data to justify postulating probability distributions over large landscapes (for discussion, see SA D.3), the standard biological intuition is that natural landscapes are at least a bit rugged and have multiple peaks. The NK-model is a family of fitness landscapes [29,30] that was introduced to study this ruggedness. This model allows tuning the amount of epistasis: the fitness contribution of each of the *n* loci depends not only on its gene, but also on the genes at up to *K* other loci (SA Definition 25).

For *K ≥* 2, the NK-model can generate hard fitness landscapes where from some initial genotypes, any adaptive walk to any local peak is exponentially long (SA Corollary 28). On such landscapes, any adaptive evolutionary dynamic – including, but not limited to, all the SSWM dynamics we’ve considered so far – generally requires an exponential number of steps to reach a local fitness optimum. Even if an omniscient navigator could always choose the most clever adaptive single mutation to arise, the adaptive path would be unbounded over polynomial timescales.

To better integrate the numeric structure of fitness, let us consider a genotype *x* to be at an *s*-approximate peak [31] if each of *x*’s mutational neighbours *y* have fitness *w*(*y*) *≤* (1 + *s*)*w*(*x*) (SA Definition 31). On the hard rough fitness landscapes described above, fittest mutant dynamics will encounter an *s*-approximate peak with moderately small *s* in a moderate number of mutational steps (polynomial in *n* and 1*/s*; SA Theorem 33).

However, on hard fitness landscapes, it is not possible to find an *s*-approximate peak for very small *s* in a feasible amount of time (i.e. not possible in time polynomial in *n* and ln 1*/s*; SA Theorem 35). This (un)reachability of *s*-approximate fitness peaks is especially important to consider in discussions of nearly-neutral networks and approximate fit-ness plateaus [7,32]. In an idealized, unstructured population, we can expect random drift to overcome selection when *s* drops below about 1*/P* where *P* is the number of individuals in the population. But certain structured populations can act as amplifiers of selection [33] and prevent drift from dominating until *s* is significantly closer to zero.

Given that the quantity *s* in the definition of an *s*-approximate peak is defined in the same way as the selection coefficient of population genetics [34], the above approximation results allow us to link the distinction between easy and hard fitness landscapes to the rich empirical literature on fitness traces and declining fitness gains in microbial evolution experiments [35]. On the hard rugged fitness landscapes described above – and even on the winding semi-smooth landscape of Figure 3 and SA C.2 – this selective coefficient drops off at the slow rate of *s*(*t*) *≈* 1*/t* for fittest mutant dynamics. In general, on any family of landscapes – even the hardest ones – *s*(*t*) can decay as fast as a power law. On easy landscapes, it can decay faster. But the power law decay in selection coefficient is the fastest decay possible on hard fitness landscapes. In particular, the selective coefficient, on hard landscapes, cannot decrease at the exponential rate (i.e. *s*(*t*) *≈ e*^*-t*^; SA Corollary 36) that is typical of equilibration in non-biological systems. This slow decay in selection coefficient is consistent with the rule of declining adaptability observed in various microbial long-term evolution experiments [35–37], suggesting that at least some naturally occurring microbial fitness landscapes might be hard. Thus, a natural candidate for hard landscapes might be the landscapes with unbounded growth in fitness observed in the *E. Coli* long-term evolutionary experiment [36]. Whereas when one sees a power-law in allometry, one expects potential physical constraints; I propose that when one see a power-law in selection strength or fitness, one should look for a computational constraint.

The existence of hard landscapes allows us to explain openended evolution as a consequence of the ultimate constraints of computational complexity. I am certainly not the first to note that populations might undergo unbounded increases in fitness and open-ended evolution. In fact, there is an extensive literature on the rate of adaptation [35, 36, 38– 42] that seems to assume (at least implicitly) that a fitter mutation is always available. These models often directly built-in unbounded growth by treating mutations as iid random samples from a distribution of fitness effects that can always generate a higher fitness variant, albeit with low probability. This approach corresponds to implicitly construction a fitness landscape as an infinite unbounded tree that lacks the second-order and higher combinatorial structure that mutation-graphs provide. These unbounded tree models are better suited to empirical operationalization than (the exponentially large but) finite fitness landscapes and make good effective theories on shorter timescale (where time – measured in the number of fixations – is significantly less than the size of the genome). But these models assume (often by reference to recent environmental change) that a beneficial mutation is always possible, rather than explain why such a mutation is always possible.

This is in stark contrast to the work I present here. To avoid building in the unbounded growth in fitness that I aim to explain, I consider families of finite fitness landscapes. I showed that these can be either easy or hard. In the hard families of landscapes, there is a computational constraint on evolution that ensures that beneficial mutations are available for effectively ever. Thus, this work can be read as an explanatory compliment to the unbounded tree models. Of course, given that I consider large but finite fitness landscapes, it is conceivable that a population will be found at a local fitness peak of a hard fitness landscape. This is conceivable in the same way as – according to the Poincare recurrence theorem – all the oxygen molecules in a large room will eventually return arbitrarily close to the corner they were released from. But just as the Poincare recurrence theorem does not invalidate the second law of thermodynamics [43], the existience of local peaks in finite static landscapes does not invalidate open-ended evolution on hard fitness landscapes.

### Arbitrary evolutionary dynamics: learning & cooperation

As we move from single genes [27, 28], to microbes [35, 36], and on to large organisms, a richer space of possible evolutionary dynamics opens up. To capture this rich space of possibilities, we need to abstract beyond adaptive dynamics by considering arbitrary mutation operators, demographies, population structures and selection functions – even ones that can cross fitness valleys and distribute the population over many genotypes. From the perspective of constraints on evolution: I want to relax the selective constraint that confines populations to an adaptive path ([44]; SA Defintion 1). By allowing non-adaptive changes, I want to highlight the power of the constraint of computational complexity, even in the absence of the selective constraint. From the perspective of evolutionary forces: we have to allow for strong forces that can potentially overpower or boost natural selection. To make sure that we have considered all possibilities, I will model arbitrary evolutionary dynamics as all polynomial-time algorithms. This takes us into the realm of the computational complexity class of polynomial local search (PLS; Johnson, Papadimitriou, and Yannakakis [45], Roughgarden [46] and SA D). But even for these most permissive population-updating procedures, evolution will in general require an infeasible amount of time to find a local fitness peak in the NK-model with *K ≥* 2 (SA Theorem 27 and Corollary 29), or to find an *s*-approximate peak for very small *s* (SA Theorem 35). Evolution will be trapped in the mazes of hard fitness landscapes and not reach anywhere near the ‘exit’ of a local fitness optimum. No proximal cause can overpower the ultimate constraint of computational complexity.

If one is accustomed to seeing results only for particular evolutionary algorithms then the generality of the above results might seem fantastical. But these are exactly the kind of general results that are typical in computational complexity theory.

The strength of this ultimate constraint allows us to reason rigorously from disequilibrium to establish positive results. For instance, that costly learning (Baldwin effect [47,48]) can remain adaptive, or that hitchhiking can maintain cooperation (Hankshaw effect [49]) effectively forever. In the case of costly learning, Simpson [48] noted: “[c]haracters individually acquired by members of a group of organisms may eventually, under the influence of selection, be reinforced or replaced by similar hereditary character”. For Simpson [48] this possibility constituted a paradox: if learning does not enhance individual fitness at a local peak and would thus be replaced by simpler non-learning strategies then why do we observe the costly mechanism and associated errors of individual learning? A similar phenomenon is important for the maintenance of cooperation. Hammarlund, Connelly, Dickinson, and Kerr [49] consider a metapopulation that is not sufficiently spatially structured to maintain cooperation. They augment the metapopulation with a number of genes with non-frequency dependent fitness effects that constitute a static fitness landscape. If adaptive mutations are available then cooperators are more likely to discover them due to the higher carrying capacity of cooperative clusters. This allows cooperation to be maintained by hitchhiking on the genes of the static fitness landscape. Hammarlund, Connelly, Dickinson, and Kerr [49] call this hitchhiking the Hankshaw effect and for them it constitutes a transient: since cooperation does not enhance opportunities for adaptive mutations at the fitness peak, then cooperators will be out-competed by defectors.

Currently, both the Baldwin and Hankshaw puzzles are resolved in the same way: just in time environmental change. Most resolutions of the Baldwin paradox focus on non-static fitness in rapidly fluctuating environments that are compatible with the speed of learning but not with evolutionary adaptation. Similarly, Hammarlund, Connelly, Dickinson, and Kerr [49] suggest making their transient permanent by focusing on dynamically changing environments. But, these just-in-time dynamic changes in the fitness landscape are not necessary if we acknowledge the existence of hard static fitness landscapes. Individual costly learning and higher densities of cooperative clusters leading to more mutational opportunities are two very different evolutionary mechanisms for increased adaptability. But they are both just polynomial time algorithms. Regardless of how much these mechanisms speed-up, slow-down, guide, or hinder natural selection, the population will still not be able to find a local fitness optimum in hard fitness landscapes. Without arriving at a fitness optimum, the paradox of costly learning disolves and the Hankshaw effect can allow for perpetual cooperation. This suggests that if we want a family of natural examples of evolution on hard fitness landscapes among more complex organisms then good candidates might be populations with costly learning or persistent cooperation. More generally, the non-vanishing supply of beneficial mutations on hard landscapes can allow selection to act on various mechanisms for evolvability [44] by letting the evolvability modifier alleles hitchhike on the favourable alleles that they produce.

These examples can be seen as instances of a more general observation on adaptationism. It is standard to frame adaptationism as “the claim that natural selection is the only important cause of the evolution of most nonmolecular traits and that these traits are locally optimal” [50]. Here, I showed that these are two independent claims. Even if we assume that (1) natural selection is the dominant cause of evolution then – on hard fitness landscapes – it does not follow that (2) traits will be locally optimal. Given the popularity of equilibrium assumptions in evolutionary biology, I expect that a number of other paradoxes and effects in addition to the Baldwin and Hankshaw could be eased by recognizing the independence of these two claims.

For those biologists who have moved on from debates about adaptationism and instead aim to explain the relative contribution of various evolutionary forces to natural patterns, I provide a new consideration: hard landscapes allow the force of natural selection on its own to explain patterns such as, for example, maladaptation. Prior accounts of maladaption rely on forces like deleterious mutation pressure, lack of genotypic variation, drift and inbreeding, and gene flow acting opposite natural selection resulting in a net zero force and thus a maladaptive equilibrium away from a fitness peak [2]. The ultimate constraint of computational complexity allows for perpetual maladaptive disequilibrium even in the absence of (or working against) these other forces.

Currently, finding a species away from a local fitness peak is taken as motivation for further questions on what mechanisms or non-selective evolutionary forces cause this discrepancy. In this context, my results provide a general answer: hard landscapes allow adaptationist accounts for the absence of evolutionary equilibrium and maladaptation even in experimental models with static environments – and/or the abscence of strong evolutionary forces working against natural selection – like the tRNA gene in yeast [28,51] or the long-term evolutionary experiment in *E. coli* [36]. By treating evolution as an algorithm, we see that time can be a limiting resource even on evolutionary timescales. These hard landscapes can be finite and deceptively simple – having only limited local epistasis or not having reciprocal sign-epistasis – and yet allow for unbounded fitness growth.

In contrast, a system found at a local fitness peak – like the snoRNA gene in yeast [27] – currently merits no further questions. The results in this report show that establishing evolutionary equilibrium should not be the end of the story. We need to also explain what features of the relevant fitness landscapes make them easy: i.e. explain why these fitness landscapes do not produce a computational constraint on evolution. For this, the tools of theoretical computer science can be used to refine our logical characterization of such fitness landscapes to guarantee that local peaks can be found in polynomial time. For example, we could consider limits on the topology of gene-interaction network (SA D.1), or the type of interaction possible between genes [52] to separate easy from hard landscapes. This opens new avenues for both empirical and theoretical work.

In this report, I provided mathematical constructions for hard fitness landscapes and suggested some empirical candidates. By doing this, I showed that computational complexity is an ultimate constraint on (our models of) evolution. But I did not establish that it is a *major* constraint in nature. Although given the empirical candidates that I suggested, I expect it to play a major role. However, after future empirical investigations, it could be that we find no naturally occurring hard fitness landscapes. This would not be a disappointment. If our models of fitness landscapes allow for ultimate constraints but we do not see those ultimate constraints in nature then we will know the direction in which to refine our models.

Given the limited – albeit growing [17, 20, 23–25, 27, 28, 51, 53, 54] – empirical data available on the distribution of natural fitness landscapes, it is tempting to turn to theoretical distributions of fitness landscapes. But we should be cautious here. The popular uniform distributions over gene-interaction networks and interaction components was introduced for ease of analysis rather than some foundational reason or empirical justification. With this distribution, I would expect hard instances to be scarce based on arguments similar to Tovey [55] and Hwang, Schmiegelt, Ferretti, and Krug [8]. However, instead of choosing a distribution for ease of analysis, we could instead choose one by Occam’s razor: i.e. the Kolmogorov universal distribution (sampling landscapes with negative log probability proportional to their minimum description length). In the Occam case, I would expect the fraction of fitness landscapes that are hard to be significant based on results similar to Li and Vitányi [56]. I leave it as an open question for future work to determine what choices of distribution of fitness landscapes are most appropriate, and how average case analysis over those particular distributions compares to the distribution-free analysis that I presented here (see SA D.3 for more discussion).

On easy landscapes, it is reasonable to assume that evolution finds locally-well-adapted genotypes or phenotypes. We can continue to reason from fitness peaks, debate questions of crossing fitness valleys, and seek solutions to Wright [1]’s problem of “a mechanism by which the species may continually find its way from lower to higher [local] peaks”. But with hard landscapes, it is better to think of evolution as open ended and unbounded. We will have to switch to a language of “adapting” rather than “adapted”. We will have to stop reasoning from equilibrium – as I did in the discussion of maintaining costly learning and cooperation. Finally, we will have to stop asking about the basins of attraction for local peaks and instead seek mechanisms that select which unbounded adaptive path evolution will follow. It is tempting to read this language of disequilibrium and negation of “locally adapted” as saying that organismal traits are not well honed to their environment. But we must resist this mistake and we must not let better be the enemy of good. Finding local optima in the hardest landscapes is a hard problem for any algorithm, not just biological evolution. In particular, it is also hard for scientists: on hard landscapes we cannot find optimal solutions either, and so the adapting answers of evolution can still seem marvelously well honed to us. And although I have focused on biological evolution, we can also look for hard landscapes in other fields. For example, these results translate directly to areas like business operation & innovation theory, where the NK-model is used explicitly [57, 58]. In physics, the correspondence between spin-glasses and the NK-model can let us look at energy minimization landscapes. In economics, classes of hard fixed-point problems similar to PLS are a lens on markets [46]. In all these cases, theoretical computer science and combinatorial optimization offer us the tools to make rigorous the distinction between easy and hard landscapes. They allow us to imagine hard landscapes not as low-dimensional mountain ranges but as high-dimensional mazes that we can search for effectively ever.

## Acknowledgements

I am grateful to Julian Xue for introducing me to the NK-model and many helpful discussions; and to Peter Jeavons for extensive feedback and encouragement. This paper benefited from comments by Eric Bolo, Frederic Guichard, Marc Harper, Sergey Kryazhimskiy, Andriy Marusyk, Daniel Nichol, Prakash Panangaden, Joshua Plotkin, Jacob Scott, participants of the 2014 Computational Theories of Evolution workshop at the Simons Institute for the Theory of Computing, and the reviewers. This work was started while I was at the School of Computer Science, McGill University.

## Appendices

In the main text, I focused on the biological importance, interpretation, and implication of these results. In these appendices, I provide the formal proof of the results. Below, I formally define the concepts introduced within the body of the report and prove the theorems on which the conclusions are based. Some of this was first presented in the Kaznatcheev [13] preprint. The structure of the appendices is below:

A Formal definitions of fitness functions, fitness landscapes, fitness graphs, and adaptive paths. I focus specific attention on epistasis (A.1) because it can be used to define broad families of landscapes, such as:

B *Smooth fitness landscapes*: these are the source of a lot of intuition and early models of fitness landscapes. So, I briefly remind the reader of important properties of smooth landscapes.

C *Semi-smooth fitness landscapes*: these share many properties in common with smooth fitness landscapes, and I prove a characterization Theorem 14 that is structured in a similar way to smooth landscapes. However, computationally, semi-smooth landscapes, unlike smooth ones, can be hard. In subsection C.1, I use the equivalence of semi-smooth fitness landscapes and acyclic unique-sink orientations of hyper-cubes to adapt hardness results from the analysis of simplex algorithms. This provides hard landscapes for fitter mutant SSWM dynamics. In the subsequent subsections, I show how to construct hard fitness landscapes for fittest mutant SSWM dynamics from specific start position (C.2) and random start position (C.3).

D *NK-model of fitness landscapes*: this is a tunable rugged fitness landscape model that – unlike the previous two – can have many peaks. To analyze this model of landscapes, I review the complexity class PLS, show that the NK-model is PLS complete for *K* ≥ 2, and discuss the generality of the results. In subsection D.1, I focus on easy instances of the NK-model and, in subsection D.3, provide an intuition for why the assumption of simple distributions of fitness landscapes in prior work might have made the existence of hard families more difficult to spot earlier. In subsection D.2, I discuss nearly-neutral networks and the hardness of *s*-approximate peaks (Definition 31).

To recap, I argue that local fitness optima may not be reachable in a reasonable amount of time even when allowing progressively more general and abstract evolutionary dynamics. For this generality, we pay with increasing complication in the corresponding fitness landscapes. This progression of results is summarized in Table 1 (which also serves as a guide for navigating the appendix). If we restrict our evolutionary dynamics to fitter or fittest mutant SSWM, then just sign epistasis is sufficient to ensure the existence of hard landscapes. If we allow any *adaptive* evolutionary dynamics, then reciprocal sign epistasis in the NK model with *K* ≥ 2 is sufficient for hard landscapes. If we want to show that arbitrary evolutionary dynamics cannot find local fitness optima then we need *K* ≥ 2 and the standard conjecture from computational complexity that FP ≠ PLS.

### A Fitness landscapes, graphs, and adaptive paths

In 1932, Wright introduced the metaphor of a fitness landscape [1]. The landscape is a genetic space where each vertex is a possible genotype and an edge exists between two vertices if a single mutation transforms the genotype of one vertex into the other. In the case of a biallelic system we have *n* loci (positions), at each of which it is possible to have one of two alleles, thus our space is the *n*-bit binary strings {0, 1}^*n*^. We could also look at spaces over larger alphabets; for example, 4 letters for sequence space of DNA, or 20 letters for amino acids; but the biallelic system is sufficiently general for us. A mutation can flip any loci from one allele to the other, thus two strings *x, y* ∈ {0, 1}^*n*^ are adjacent if they differ in exactly one bit. Thus, the landscape is an *n*-dimensional hypercube with genotypes as vertices. The last ingredient, fitness, is given by a function that maps each string to a non-negative real number. For the majority of this report – with the exception of subsection D.2 – the exact fitness values or their physical interpretations do not matter. Only their rank-ordering matters.

Individual organisms can be thought of as inhabiting the vertices of the landscape corresponding to their genotype. And we imagine evolution as generally trying to ‘climb uphill’ on the landscape by moving to vertices of higher fitness.

**Definition 1.** In a fitness landscape with fitness *f*, a path *v*_1_*…v*_*t*_ is called *adaptive* if each *v*_*i*+1_ differs from *vi* by one bit and *f* (*v*_*i*+1_) > *f* (*v*_*i*_).

These are sometimes also called *accessible* paths, but I will avoid this terminology because for the most general evolutionary dynamics, the paths taken don’t have to be strictly increasing in fitness; i.e. they don’t have to necessarily be adaptive. In other words, for arbitrary evolutionary dynamics, evolution can follow (or access) non-adaptive paths. So non-adaptive paths are accessible to arbitrary evolutionary dynamics; and it would be awkward to say that non-accessible paths are accessible to arbitrary evolutionary dynamics. If some particular evolutionary dynamic produces only adaptive paths, though, then it is called an *adaptive dynamic*. In general, adaptive paths can continue until they reach a local fitness optimum.

**Definition 2.** A genotype *u* is a *local fitness optimum* (sometimes also called a *(local) fitness peak*) if for all adjacent genotypes *v*, we have *f* (*v*) ≤ *f* (*u*).

Note the above definition of a genotype as a local fitness optimum allows for adjacent genotypes of equal (or lesser) fitness. In particular, this means that points within a *fitness plateau* can be local fitness optima. At times, I will assume for simplicity that no two adjacent genotypes have exactly the same fitness to avoid considering fitness plateaus; but this is not an important restriction and all the hardness results can be reproved without it. A local fitness optimum is a *global fitness optimum* if all other genotypes in the whole of the genetic space (not just neighbours) have the same or lower fitness; i.e. if no other local fitness optimum in the whole of the genetic space has a higher fitness.

Sometimes it is useful to represent a fitness landscape as a *fitness graph* by replacing the fitness function by a flow: for adjacent genotypes in the mutation-graph, direct the edges from the lower to the higher fitness genotype. This results in a characterization of fitness landscapes of a biallelic system as directed acyclic graphs on {0, 1}^*n*^. Fitness peaks would correspond to sinks, and adaptive paths would correspond to paths that follow the edge directions of the DAG. I will consider a population at evolutionary equilibrium if it finds a local peak in the fitness landscape; i.e. a sink in the fitness graph. Crona, Greene, and Barlow [19] introduced this representation into theoretical biology, but fitness graphs have been used implicitly in earlier empirical studies of fitness landscapes [24, 59–61]. Using fitness graphs is particularly useful empirically because it is difficult to quantitatively compare fitnesses across experiments. However, if pairwise competitions are used to build an empricial fitness graph, it is important to verify that the graph is transitive (acyclic) [62]. In theoretical work, the fitness graph approach has made the proofs of some classical theorems relating local structure to global properties easier and shifts our attention to global algorithmic properties of evolution instead of specific numeric properties.

Throughout the report and these appendices, I consider individual genotypes or phenotypes as the domain of the fitness landscapes. Thus, I am focusing on micro-evolutionary processes. Given the extremely long time scales that I am considering in this report, it is also natural to consider generalizations where the vertices in the fitness landscape are interpreted as whole species and mutations as speciation events. For simplicity, I will not explicitly discuss such macro-evolutionary processes.

#### A.1 Epistasis

*Epistasis* is a measure of the kind and amount of inter-locus interactions. Consider two loci with the first having alleles a or A, and the second b or B. Assume that the upper-case combination is more fit: i.e. *f* (*ab*) < *f* (*AB*).

**Definition 3.** Two alleles are *non-interacting* if the fitness effects are additive and independent of background: *f* (*AB*) - *f* (*aB*) = *f* (*Ab*) - *f* (*ab*), *f* (*AB*) - *f* (*Ab*) = *f* (*aB*) - *f* (*ab*).

In *magnitude epistasis* this additivity is broken, but the signs remain: *f* (*AB*) > *f* (*aB*) > *f* (*ab*) and *f* (*AB*) > *f* (*Ab*) > *f* (*ab*). The difference between non-interacting alleles and magnitude epistasis is not invariant under rankorder preserving transformation of the fitness function, thus I will not distinguish between the two types. Throughout the paper, I will use ‘no epistasis’ to mean both non-interacting alleles and magnitude epistasis, as the following definition makes explicit.

**Definition 4.** If *f* (*AB*) > *f* (*aB*) > *f* (*ab*) and *f* (*AB*) > *f* (*Ab*) > *f* (*ab*) then we will say that there is *no epistasis* between those alleles.

A system has sign epistasis if it violates one of the two conditions for magnitude epistasis. For example, if the second locus is b then the mutation from a to A is not adaptive, but if the second locus is B then the mutation from a to A is adaptive.

**Definition 5.** Given two loci, if *f* (*AB*) > *f* (*aB*) > *f* (*ab*) > *f* (*Ab*) then there is *sign epistasis* at the first locus.

Finally, a system has reciprocal sign epistasis if both conditions of magnitude epistasis are broken, or if we have sign epistasis on both loci [17–19].

**Definition 6.** Given two loci, if *f* (*AB*) ≥ *f* (*ab*) but *f* (*ab*) > *f* (*Ab*) and *f* (*ab*) > *f* (*aB*) then there is *reciprocal sign epistasis* between those two loci.

Figure 4 visualizes all the fitness graphs on two loci and categorizes the type of epistasis present.

**Figure 4:**
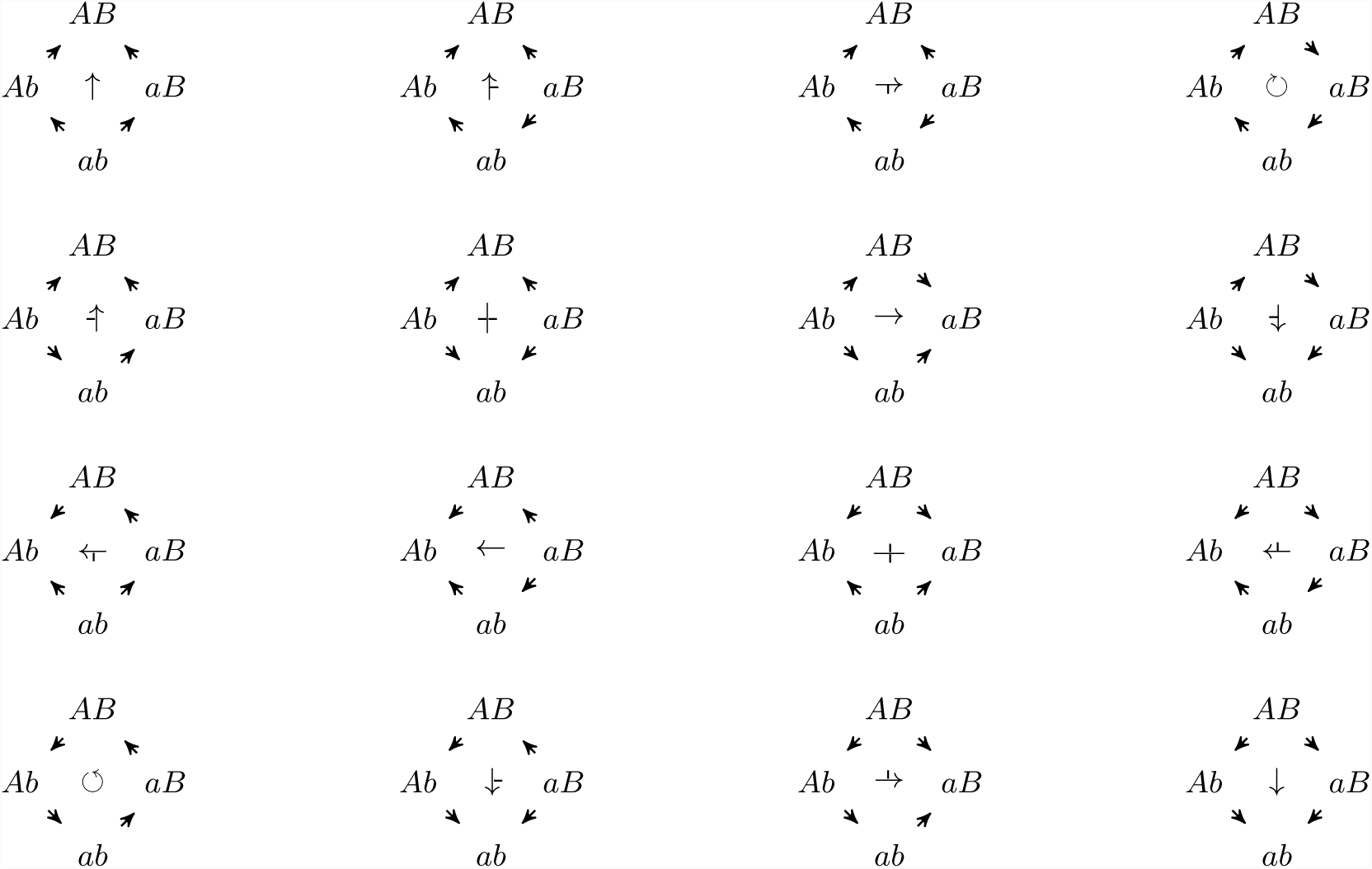
Three different kinds of epistasis possible in fitness graphs: no epistasis (↑), sign epistasis (-↑, ↑-), and reciprocal sign epistasis (†). Arrows in the fitness graph are directed from lower fitness genotypes towards mutationally adjacent higher fitness genotypes. In the middle of each fitness graph is a symbol showing the kind (and orientation) of epistasis. Note that the bottom left (↺) and top right (↻) fitness graphs violate transitivity.

### B Smooth fitness landscapes

If a fitness landscape has no sign epistasis then it is a smooth landscape and has a single peak *x*^*^ [19, 21]. Every shortest path from an arbitrary *x* to *x*^*^ in the mutation-graph is an adaptive path – a flow in the fitness graph – and every adaptive path in the fitness graph is a shortest path in the mutation graph [19]. Thus, evolution can quickly find the global optimum in a smooth fitness landscape, with an adaptive path taking at most *n* steps: that is, all smooth fitness landscapes are easy landscapes. For an example, see the smooth *Escherichia coli β*-lactamase fitness landscape measured by Chou, Chiu, Delaney, Segre, and Marx [20] in Figure 2a.

**Proposition 7** ([19,21]). *If there is no sign epistasis in a fitness landscape, then it is called a* smooth landscape *and has a single peak x*^*^. *Every shortest path (ignoring edge directions) from an arbitrary genotype x to x*^*^ *is an adaptive path, and every adaptive path from x to x*^*^ *is a shortest path (ignoring edge directions).*

Where by ‘shortest path (ignorning edge directions)’, I mean any shortest path in the mutation-graph, irrespective of if the fitness along the edges of that path increases (‘up arrow’) or decreases (‘down arrow’). In other words, an arbitrary shortest path between *x* and *x*^*^ corresponds to an arbitrary swapping of genes at the loci on which *x* and *x*^*^ from the value they have in *x* to the value in *x*^*^. This means, for example, that if *x* and *x*^*^ differ on *d* loci then there are *d*! many shortest paths between them and by Proposition 7 those are also the *d*! adaptive paths between them.

### C Semi-smooth fitness landscapes

Since a smooth landscape is always easy, let’s introduce the minimal amount of epistasis: sign epistasis, without any reciprocal sign epistasis.

**Definition 8.** A *semi-smooth fitness landscape* on {0, 1}^*n*^ with fitness function *f* is a fitness landscape that has no reciprocal sign epistasis. Such a fitness function *f* is also called semi-smooth.

For some of the following proofs, it will be useful to define sublandscapes.

**Definition 9.** Given a landscape on *n* bits, a sublandscape spanned by *S* ⊆ [*n*] is a landscape on {0, 1}^*S*^ where the alleles at the loci (indices) in *S* can vary but the indices in [*n*] - *S* are fixed according to some string *u* ∈ {0, 1}^[*n*]-*S*^.

Note that the whole landscape is a sublandscape of itself (taking *S* = [*n*]). For any *S* ⊂ [*n*], there are 2^*n*-|*S*|^ many sublandscapes on *S* corresponding to the possible *u* ∈ {0, 1}^[*n*]-*S*^. Reciprocal sign epistasis between bits *i* and *j* corresponds to a sublandscape on {*i, j*} that has two distinct peaks.

Now, I can note a couple of important properties of semi-smooth landscapes:

**Proposition 10.** *If a fitness landscape on* {0, 1}^*n*^ *has some sublandscape with more than one distinct peak then it has reciprocal sign epistasis.*

The proof will show that a minimal multi-peak sublandscape must have size 2. I will do this by considering longest walks in a sublandscape. The proposition is similar to the one proved by Poelwijk, Sorin, Kiviet, and Tans [18], although my proof is distinct.

*Proof.* Let’s consider a minimal sublandscape *L* that has more than one distinct peak: that means that if this sublandscape is spanned by *S* (i.e. {0, 1}^*S*^) then no sublandscape spanned by *T* ⊂ *S* has multiple peaks.

Since *L* is minimal, its peaks must differ from each other on each bit in *S*, for if there was a bit *i* ∈ *S* on which two peaks agreed then that bit could be fixed to that value and eliminated from *S* to make a smaller sublandscape spanned by *S* -{*i*} with two peaks. Thus, the minimal multipeak sublandscape has precisely two peaks. Call these peaks *x*\* and *y*\*.

**Claim:** In a minimal multipeak sublandscape, from each non-peak vertex, there must be a path to each peak.

Let’s prove the claim by contradiction: Consider an arbitrary non-peak vertex *x*, and suppose it has no path to the *x*^*^ peak. Since any path from *x* in *L* must terminate at some peak, take the longest path from *x* to the peak *y*^*^ that it reaches, and let *y* be the last step in that path before the peak. Notice that *y* must only have one beneficial mutation (on bit *i*), the one to the peak. For if it had more than one beneficial mutation, it could take the non-peak step to *y′* and then proceed from *y′* to *y*^*^ (*x*^*^ is not an option by assumption, and there are only two peaks in *L*) and thus provide a longer path to the peak. Now consider the landscape on *S* -{*i*}, with the *i*th bit fixed to *y*_*i*_. Since *y*_*i*_ is the same as 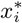 (both are opposite of 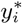), *x*^*^ is still a peak over *S* -{*i*}, but so is *y* (since it’s only beneficial mutation was eliminated by fixing *i* to *y*_*i*_). But this contradicts minimality, so no such *x* exists.

Now that we know that we can reach each peak from any vertex *x*, let us again consider the longest path from *x* to *y*^*^ with *y* as the last step in that path before the peak, and *i* as the position of the last beneficial mutation. Since all non-peak vertices must reach both peaks, there must be some other beneficial mutation *j* from *y* to *x′* that eventually leads to *x*^*^. But if *x′* is not a peak then it must also have a way to reach *y*^*^, but then we could make a longer path, contradicting the construction of *y*. Thus *x′* must be the peak *x*^*^.

This means that *x*^*^ and *y*^*^ differ in only the two bits *i* and *j*. But in a minimal multipeaked sublandscape they must differ in all bits, so *S* = {*i, j*}; i.e. this sublandscape is an example of reciprocal sign epistasis.□

**Corollary 11.** *A fitness landscape without reciprocal sign epistasis has a unique single peak.*

*Proof.* This follows from the contrapositive of Proposition 10, since the whole landscape is a sublandscape of itself.□

The above results can also be restated in the terminology used to analyze simplex algorithms.[22,26]

**Definition 12.** A directed acyclic orientation of a hypercube {0, 1}^*n*^ is called an *acyclic unique sink orientation (AUSO)* if every subcube (face; including the whole cube) has a unique sink.

This makes the contrapositive of Proposition 10 into the following proposition:

**Proposition 13.** *A semi-smooth fitness landscape is an AUSO*

Now, if we let *x* ⊕ *y* mean XOR between *x* and *y* and let ‖*z*‖_1_ mean the number of 1s in *z* then we can state the main theorem about semi-smooth fitness landscapes:

**Theorem 14.**

*A semi-smooth fitness landscape has a unique fitness peak x*^*\**^ *and for any vertex x in the landscape, there exists a path of length* ‖*x*^*\**^ ⊕ *x*‖_1_ *(Hamming distance to peak) from x to the peak.*

*Proof.* The unique peak *x*^*\**^ is just a restatement of Corollary 11. To show that there is always a path of Hamming distance to the peak, I will show that given an arbitrary *x*, we can always pick a mutation *k* that decreases the Hamming distance to *x*^*\**^ by 1.

Let *S* be the set of indices that *x* and *x*^*\**^ disagree on, |*S*| = ‖*x*^*\**^ ⊕ *x*‖1. Consider the sub-landscape on *S* with the other bits fixed to what *x* and *x* agree on. In this sublandscape *x* is a peak, thus by Proposition 10 *x* isn’t a peak and must have some beneficial mutation *k ∈ S*. This is the *k* we were looking for.□

Note that this proof specifies an algorithm for constructing a short adaptive walk to the fitness peak *x*^*\**^. However, this algorithm requires knowing *x*^*\**^ ahead of time – i.e. seeing the peak in the distance. But evolution does not know ahead of time where peaks are, and so cannot carry out this algorithm. Even though a short path to the peak always exists, evolutionary dynamics might not follow it.

#### C.1 Hard landscapes for random fitter SSWM

The simplest evolutionary rule to consider is picking a mutation uniformly at random among ones that increase fitness. This can be restated as picking and following one of the out-edges in the fitness graph at random; i.e. this is equivalent to the random-edge simplex pivot rule [26]. Proposition 13 allows me to use the hard AUSOs constructed by Matousek and Szabo [26] as a family of hard semi-smooth landscapes.

**Theorem 15** ([26] in biological terminology). *There exist semi-smooth fitness landscapes on* {0, 1}^*n*^ *such that random fitter mutant SSWM dynamics starting from a random vertex, with probability at least* 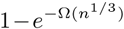 *follows an adaptive path of at least* 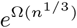 *steps to evolutionary equilibrium.*

In other words, multiple peaks – or even reciprocal sign-epistasis – are not required to make a complex fitness landscape. In fact, AUSOs were developed to capture the idea of a linear function on a polytope (although AUSOs are a slightly bigger class). It is not surprising to find the simplex algorithm in the context of semi-smooth landscapes, since we can regard it as a local search algorithm for linear programming where local optimality coincides with global optimality. Linear fitness functions are usually considered to be some of the simplest landscapes by theoretical biologists; showing that adaptation is hard on these landscapes (or ones like them) is a surprising result.

#### C.2 Construction of hard semi-smooth landscapes for fittest SSWM

One might object to taking random fitter mutants because sometimes the selected mutations are only marginally fitter than the wildtype. It might seem natural to speed-up evolution by always selecting the fittest possible mutant. Here I show that, in general, this does not help.

Consider a fitness landscape on {0, 1}^*m*^ with semi-smooth fitness function *f* that if started at 0^*m*^ will take *k* steps to reach its evolutionary equilibrium at *x*^*\**^. I will show how to grow this into a fitness landscape on {0, 1 }^*m*+2^ with semi-smooth fitness function *f ′* that if started at 0^*m*+2^ will take 2(*k* + 1) steps to reach its evolutionary equilibrium at 0^*m*^11.

For simplicitly of analysis, let us define the following functions and variables for all points in {0, 1}^*m*^ that aren’t an evolutionary equilibrium under *f*; i.e. all except *x*. Let

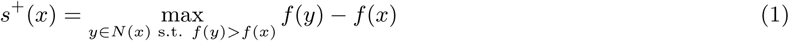

and

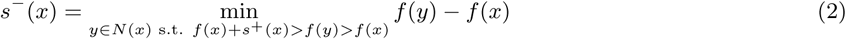

where *N* (*x*) are the neighbours of *x* in the mutation graphs; i.e. genotypes that differ from *x* in one bit.

Now overload these into constants, as follows: define *s*^+^ = min_*x*_ *s*^+^(*x*) and *s*^−^ = min*x s*^−^(*x*). Suppose that *f* is such that *s*^−^ < *s*^+^; otherwise set *s*^−^ = *s*^+^*/*2 (do this also, if *N* (*x*) s.t. *f* (*x*) + *s*^+^(*x*) > *f* (*y*) > *f* (*x*) is empty for some non-equilibrium *x*).

Let *x* ⊕ *y* mean the XOR between *x* and *y*. Consider the ‘reflected’ function *f* (*x* ⊕ *x*^*\**^). I call this function *reflected* because we could visualize *f* (*x* ⊕ *x*^*\**^) as the same function as *f* (*x*), except it is mirrored (swapping the fitness effects of the 0 and 1 allele) along each locus *i* where 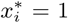. Note that if *f* (*x*) is semi-smooth then so is *f* (*x* ⊕ *x*^*\**^), since it just relabels the directions of some dimensions. The reflected function preserves all the important structure. In particular, if under *f* (*x*) it took *k* steps to go from 0^*m*^ to *x*^*\**^ then under *f* (*x ⊕x*^*\**^) it will take *k* steps to from from *x*^*\**^ to 0^*m*^.

Now define *f ′*: {0, 1}^*m*+2^ *?*ℝ as:

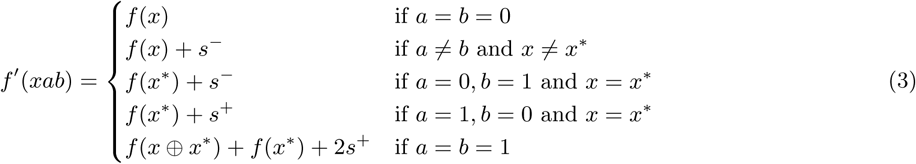

Basically the *x*00 subcube is the original landscape, the *x*10 and *x*01 subcubes serve as ‘buffers’ to make sure that the walk doesn’t leave the first subcube before reaching *x*^*\**^00, and the *x*11 is the original landscape reflected around *x*^*\**^ that takes us from *x*^*\**^11 to 0^*m*^11.

Notice, that *f ′* has the same *s*^+^ and *s*^−^ as *f*.

Now we just need to establish some properties:

**Proposition 16.** *Fittest mutant SSWM dynamics will not leave the* {0, 1}^*m*^00 *subcube until reaching x*^*\**^00.

*Proof.* By definition, the fittest mutant (i.e. neighbour over {0, 1}^*m*^) from each genotype *x ∈* {0, 1}^*m*^ that isn’t *x*^*\**^ in *f*, has a fitness advantage of *s* or higher. Hence adding two extra edges from *x*00 to *x*10 and *x*01, each with fitness advantage *s*^−^ < *s*^+^ will not change the edge that fittest-mutant SSWM picks.□

**Proposition 17.** *SSWM dynamics will not leave the* {0, 1}^*m*^11 *subcube after entering it.*

*Proof.* This is because *f ′* has strictly greater fitness on the {0, 1}^*m*^11 subcube than on the other three subcubes. Confirming this, note that for every *x ∈* {0, 1}^*m*^:

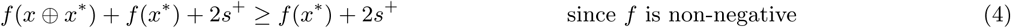

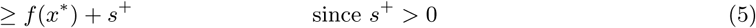

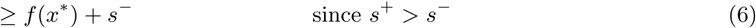

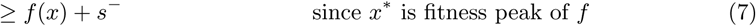

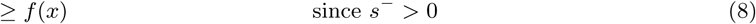

□

**Proposition 18.** *If f on* {0, 1}^*m*^ *has no reciprocal sign-epistasis then f ′ on* {0, 1}^*m*+2^ *has no reciprocal sign-epistasis.*

*Proof.* Consider any pair of genes *i, j ∈* [*m*]. Among these first *m* genes, depending the last two bits, we are looking at landscapes on {0, 1}^*m*^ 00, {0, 1}^*m*^ 01, {0, 1}^*m*^ 10, or {0, 1}^*m*^ 11, with the fitness given by *f* (*x*),*f* (*x*) + *s*^−^, *f* (*x*) + *s*^−^, or *f* (*x* ⊕ *x*^*\**^) + *f* (*x*^*\**^) + 2*s*^+^ (respectively). All these landscapes have isomorphic combinatorial structure to *f* and thus the same kinds of epistasis. Since *f* has no reciprocal sign-epistasis, all these subcubes lack it, too.

Now, let’s look at the case of where the gene pair goes outside the first *m* genes. Consider an arbitrary gene *i ∈* [*m*], let *u ∈* {0, 1}^*i-*1^, *v ∈* {0, 1}^*m-i*^ be arbitrary. Label *a, A ∈* {0, 1} such that *f* (*uav*) < *f* (*uAv*). look at the subcube *u*{0, 1}*v*{0, 1}^2^:

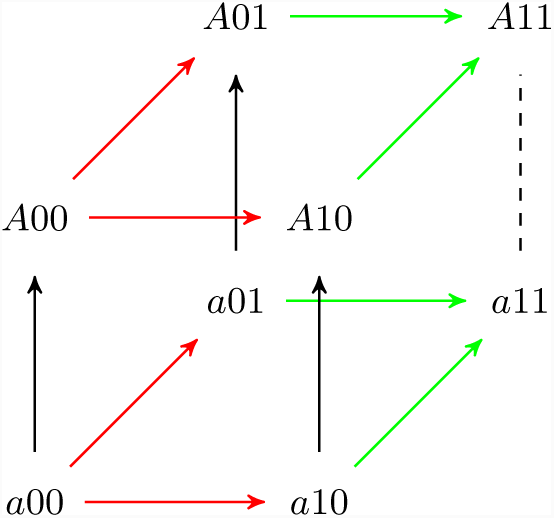

The solid black edges have their directions from the definition of *a* and *A*. The red edges have their direction because *s*^+^ > *s*^−^ > 0. The green edges have their direction because of Prop. 17. The direction of the dotted black edge will depend on if *x*^*\**^ contains 0 (point up) or 1 (point down) at position *i*, but regardless of the direction, no reciprocal sign epistasis is introduced.□

**Corollary 19.** *Given f ′ on* {0, 1} ^*m*+2^, *the fittest mutant SSWM dynamics starting at* 0^*m*+2^ *will take* 2(*k* + 1) *steps to reach its unique fitness peak at* 0^*m*^11.

*Proof.* By Prop. 16, the walk will first proceed to *x*^*\**^00 taking *k* steps. From *x*^*\**^00, there are only two adaptive mutations *x*^*\**^10 or *x*^*\**^01, and the first is fitter. From *x*^*\**^10 there is only a single adaptive mutation (to *x*^*\**^11), taking us to *k* + 2 steps. From *x*^*\**^11, by Prop. 17, it will take us *k* more steps to reach 0^*m*^11; totaling 2(*k* + 1) steps.□

**Theorem 20.** *There exist semi-smooth fitness landscapes on* 2*n loci that take* 2^*n*+1^ −2 *fittest mutant steps to reach their unique fitness peak at* 0^2(*n*-^1^)^11 *when starting from* 0^2*n*^.

*Proof.* We will build the family of landscapes inductively using our construction, starting from an initial landscape:

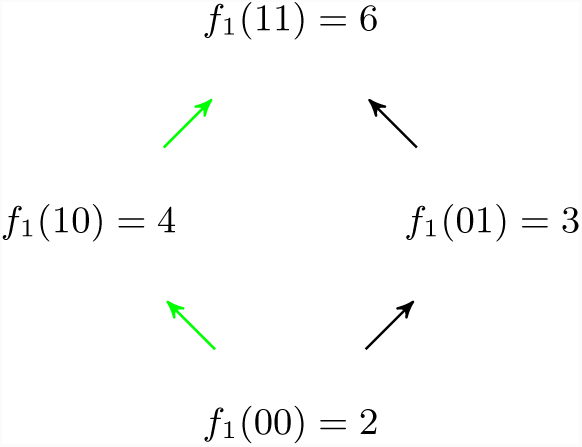

The resulting path length *T*_*n*_ will be given by the recurrence equation: *T*_*n*+1_ = 2*T*_*n*_ + 2 with *T*_1_ = 2. This recurrence is solved by *T*_*n*_ = 2^*n*+1^ −2.□

Call the landscapes constructed as in the above proof, a *winding landscapes*. A visual example of the winding landscape construction on 6 loci (*n* = 3 in Theorem 20) is given in Figure 3. The winding landscapes construction is similar to Horn, Goldberg, and Deb [63]’s *Root2path* construction, except their approach introduced reciprocal sign epistasis despite having a single peak.

Of course, this is an arbitrary initial fitness landscape and any semi-smooth landscape can be used as a starting point; the walk would still scale exponentially, but there would be a different initial condition. Further, this winding product construction I showed above is just one example for building families. Many more could be considered.

In particular, if we are interested in larger mutation operators like *k*-point mutations instead of just 1-point mutations then it is relatively straightforward to modify the winding landscape construction. As written, equation 3 uses a buffer of 2 bits in *f* ^*′*^(*xab*) to transition from *f* (*x*) to its reflection *f* (*x* ⊕ *x*^*\**^). In the more general setting, we’d pad the buffer to be *k* + 1 bits: define *f* ^*′*^(*xy*) where | *y*| = *k* + 1 with a smooth landscape on the *y* portion of the input taking us from *f* (*x*) to its reflection. Which leaves most of the above arguments unchanged, only modifying Theorem 20 to have the landscape to be on *kn* loci and the recurrence relation at the end of the proof to be *T*_*n*+1_ = 2*T*_*n*_ + *k* + 1.

#### C.3 Hard landscapes from random start

Unfortunately, one might not be impressed by a result that requires starting from a specific genotype like 0^*m*^ and ask for the expected length of the walk starting from a random vertex. Of course, if a genotype on this long walk is chosen as a starting point then the walk will still be long in most cases. However, there are only 2^*n*+1^-2 vertices in the walk, among 2^2*n*^ vertices total, so the probability of landing on the walk is exponentially small. Instead, I will rely on direct sums of landscapes and Proposition 16 to get long expected walks.

**Proposition 21.** *With probability* 1*/*4, *a winding landscape on* 2*n loci will take* 2^*n*^ *or more fittest mutant steps to reach the fitness peak from a starting genotype sampled uniformly at random.*

*Proof.* With probability 1*/*4, the randomly sampled starting vertex has the form *x*00 (i.e. its last two bits are 0s). By prop. 16, the walk can’t leave the {0, 1}^2(*n-*1)^00 landscape until reaching its peak at 0^2(*n-*2)^1100. This might happen quickly, or it might even already be at that peak. But after, it has to follow the two steps to 0^2(*n-*2)^1111 and then due to prop. 17 it will have to follow the normal long path, taking 2^*n*^ *-* 2 more steps.□

Because of the constant probability of an exponentially long walk, we can get a big lower bound on the expected walk time:

**Corollary 22.** *Fittest mutant dynamics starting from a uniformly random genotype will have an expected walk length greater than* 2^*n-*2^ *on a* 2*n-loci winding landscape.*

*Proof.* With probability 1*/*4, the the walk takes 2^*n*^ or more steps, and with probability 3*/*4 it takes 0 or more steps. Thus the expected walk length is greater than or equal to (1*/*4) *∗* 2^*n*^ + (3*/*4) *∗* 0 = 2^*n-*2^.□

However, 75% of the time, we can’t make a guarantee of long dynamics. We can overcome this limitation by taking direct sums of landscapes.

**Definition 23.** Given two fitness landscapes, one with fitness *f*_1_ on 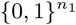 and the other with fitness 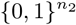, the *direct sum* (*f*_1_ *⊕ f*_2_) is a landscape with fitness 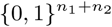 where *f* (*xy*) = *f*_1_(*x*) + *f*_2_(*y*).

Now, for any probability of failure 0 *< δ <* 1, let 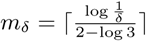 (where log is base 2).

**Theorem 24.** *There exist semi-smooth fitness landscapes on* 2*nm*_*δ*_ *loci that with probability* 1 *- δ, take* 2^*n*^ *or more fittest mutant steps to reach their fitness peak from a starting genotype sampled uniformly at random.*

*Proof.* Consider a landscape that is the direct sum of *m*_*δ*_ separate 2*n*-loci winding landscapes. Since each constituent is semi-smooth and since sums don’t introduce epistasis, the resulting ‘tensor sum’ landscape is also semi-smooth. Further, to reach its single peak, the walk has to reach the peak of each of the *m*_*δ*_ independent winding sublandscapes. But as long as at least one sublandscape has a long walk, we are happy. By prop. 21, we know that for each sublandscape, we will have a short-walk starting genotype with probability less than 3*/*4. The probability that none of them get a long walk then is less than 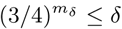.□

### D NK model with *K ≥* 2 **is PLS-complete**

**Definition 25** ([29, 30, 64]). The *NK model* is a fitness landscape on {0, 1}_*n*_. The *n* loci are arranged in a *gene-interaction network* where each locus *x*_*i*_ is linked to *K* other loci 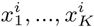 and has an associated fitness contribution function *f*_*i*_: {0, 1}^*K*+1^ *→*ℝ_+_ Given a vertex *v ∈* {0, 1}_*n*_, we define the fitness 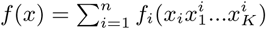.

By varying *K* we can control the amount of epistasis in the landscape. The model also provides an upper bound of 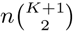 on the number of gene pairs that have epistatic interactions. Typically in the biology and statistical physics literature, the fitness contributions *f*_*i*_ and sometimes the gene-interaction network are chosen uniformly at random from some convenient probability distribution. In contrast, the approach of theoretical computer scientists is to consider *arbitrary* rather than random choices for each *f*_*i*_ and sometimes the gene-interaction network. This is an important cultural difference in methodology between statistical physics and computer science that I discuss in more detail in subsection D.3.

Weinberger [65] showed that checking if the global optimum in an *NK* model is greater than some input value *V* is *NP* -complete for *K ≥* 3. Although this implies that finding a global optimum is difficult, it says nothing about local optima. As such, it has generated little interest among biologists, although it spurred interest as a model in the evolutionary algorithms literature, leading to a refined proof of *NP* -completeness for *K ≥* 2 [66].

To understand the difficulty of finding items with some local property like being an equilibrium, Johnson, Papadimitrio & Yannakakis [45] defined the complexity class of polynomial local search (PLS). A problem is in PLS if it can be specified by three polynomial time algorithms [46]:

1. An algorithm *I* that accepts an instance (like a description of a fitness landscape) and outputs a first candidate to consider (the initial genotype).
2. An algorithm *F* that accepts an instance and a candidate and returns a objective function value (i.e. computes the fitness).
3. An algorithm *M* that accepts an instance and a candidate and returns an output with a strictly higher objective function value, or says that the candidate is a local maximum.

We consider a PLS problem solved if an algorithm can output a locally optimal solution for every instance. This algorithm does not necessarily have to use *I, F*, or *M* or follow adaptive paths. For instance, it can try to uncover hidden structure from the description of the landscape. A classical example would be the ellipsoid method for linear programming. The hardest problems in PLS – i.e. ones for which a polynomial time solution could be converted to a solution for any other PLS problem – are called PLS-complete. It is believed that PLS-complete problems are not solvable in polynomial time (i.e. FP *≠* PLS; where FP stands for the set of function problems solvable in polynomial time), but – much like the famous P *≠* NP question – this conjecture remains open. Note that finding local optima on fitness landscapes is an example of a PLS problem, where *I* is your method for choosing the initial genotype, *F* is the fitness function, and *M* computes an individual adaptive step.

**Definition 26** (Weighted 2SAT). Consider *n* variables *x* = *x*_1_*…x*_*n*_ *∈* {0, 1}_*n*_ and *m* clauses *C*_1_, *…, C*_*m*_ and associated positive integer weights *c*_1_, *…c*_*m*_. Each clause *C*_*k*_ contains two literals (a literal is a variable *x*_*i*_ or its negation 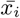), and contributes *c*_*k*_ to the fitness if at least one of the literals is satisfied, and nothing if neither literal is satisfied. The total fitness *c*(*x*) is the sum of the individual contributions of the *m* clauses. Two assignments *x* and *x*^*’*^ are adjacent if there is exactly one index *i* such that 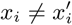. We want to maximize fitness.

The Weighted 2SAT problem is PLS-complete [67]. To show that the *NK* model is also PLS-complete, I will show how to reduce any instance of Weighted 2SAT to an instance of the *NK* model.

**Theorem 27.** *Finding a local optimum in the NK fitness landscape with K ≥* 2 *is PLS-complete.*

*Proof.* Consider an instance of Weighted 2SAT with variables *x*_1_, *…, x*_*n*_, clauses *C*_1_, *…, C*_*m*_ and positive integer costs *c*_1_, *…, c*_*m*_. We will build a landscape with *m*+*n* loci, with the first *m* labeled *b*_1_, *…, b*_*m*_ and the next *n* labeled *x*_1_, *…, x*_*n*_. Each *b*_*k*_ will correspond to a clause *C*_*k*_ that uses the variables *x*_*i*_ and *x*_*j*_ (i.e., the first literal is either *x*_*i*_ or 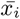 and the second is *x*_*j*_ or 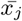; set *i < j* to avoid ambiguity). Define the corresponding fitness effect of the locus as:

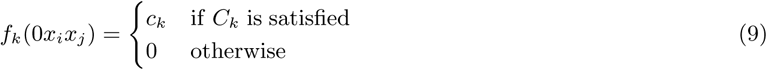

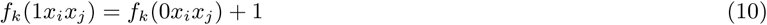

Link the *x*_*i*_ arbitrarily (say to *x*_(*i* mod *n*)+1_ and *x*_(*i*+1 mod *n*)+1_, or to nothing at all) with a fitness effect of zero, regardless of the values.

In any local maximum *bx*, we have *b* = 11..1 and *f* (*x*) = *m* + *c*(*x*). On the subcube with *b* = 11..1 Weighted 2SAT and this NK model have the same exact fitness graph structure, and so there is a bijection between their local maxima.□

Assuming – as most computer scientists do – that there exists some problem in PLS not solvable in polynomial time (i.e. FP *≠* PLS), then Theorem 27 implies that no matter what mechanistic rule evolution follows (even ones we have not discovered, yet), be it as simple as SSWM or as complicated as any polynomial time algorithm, there will be NK landscapes with *K* = 2 such that evolution will not be able to find a fitness peak efficiently. But if we focus only on rules that follow adaptive paths then we can strengthen the result:

**Corollary 28.** *There is a constant c >* 0 *such that, for infinitely many n, there are instances of NK models (with K ≥* 2*) on* {0, 1}^*n*^ *and initial genotype v such that any adaptive path from v will have to take at least* 2^*cn*^ *steps before finding a fitness peak.*

*Proof.* If the initial vertex has *s* = 11*…*1 then there is a bijection between adaptive paths in the fitness landscape and any weight-increasing path for optimizing the weighted 2SAT problem. Thus, Schaffer and Yannakakis [67]’s Theorem 5.15 applies.

This result holds independent of any complexity theoretic assumptions about the relationship between polynomial-time and PLS. Hence, there are some landscapes and initial genotypes, such that any rule we use for adaptation that only considers fitter single-gene mutants will take an exponential number of steps to find the local optimum.

If we turn to larger mutational neighbourhoods than singe-gene mutants then – due to the large class of possible adaptive dynamics – a variant of Corollary 28 will have to be reproved (often using a buffer padding argument similar to the end of section C.2) but Theorem 27 is unaffected:

**Corollary 29.** *For any definition of local equilibrium with respect to a mutation neighbourhood that contains point-mutations as a subset (i.e. if ∀x* {*y | ||y - x||*_1_ = 1} *⊆ N* (*x*)*), the NK model with K ≥* 2 *is PLS-hard.*

*Proof.* Any mutation operator that is a superset of point-mutations will only decrease the number of evolutionary equilibria without introducing new ones. Thus, it will only make the task of finding that equilibrium (just as, or) more difficult. However, since the algorithms studied by PLS do not have to use the mutation operator during their execution, changing it does not give them any more computational resources.

Finally, it is important to see the NK-model as an example model, albeit a simple and natural one. If we consider more complex models of fitness landscapes – say dynamic fitness landscapes – it is often the case that there is some parameter or limit that produces the special case of a static fitness landscape like the NK-model. In particular, static landscapes are often a sub-model of dynamic fitness landscapes and thus solving dynamic fitness landscapes can only be more difficult that static ones.

#### D.1 Easy instances of NK-model

Note that this doesn’t mean that all instances of the NK-model are hard. In fact, there are natural sub-families of the NK-model that are easy.

The simplest easy family is *K* = 0. In that case, the genes are non-interacting and we have a smooth fitness landscapes. And all smooth landscapes are easy. For *K* = 1, Wright, Thompson, and Zhang [66] presented a dynamic programming approach that can find the global fitness peak in polynomial time. Since we could use this as our algorithm *I* to pick the initial genotype, this means the model cannot be PLS-complete for *K* ≤ 1 (unless PLS = P, in which case all local search problems are easy). This means that Theorem 27 is as tight as possible in terms of *K*.

Alternatively, instead of restricting *K*, we can restrict how the gene-interaction network is connected. It will come in useful to visualize these gene-interaction networks by drawing an edge directed from a focal locus to the *K* loci that affect its fitness contribution. For example, if the genes can be arranged in a circle and a focal gene can interact with only the next *K* genes in the circle then there is a polynomial time dynamic programming algorithm to find an evolutionary equilibrium [66]. Thus, this restricted model cannot be PLS complete for any constant *K*.

It is an open question if SSWM dynamics – or some other reasonable evolutionary dynamics – is sufficient in the cases of *K* = 1 and circular arrangements. I conjecture that adaptive dynamics are sufficient in these cases, but proof of this is left for future work.

Some of the most common kinds of easy instances of the NK-model come from ‘simplifying’ restrictions put in place by the modeler. In particular, restriction on the range of the fitness function.

**Proposition 30.** *If the number of distinct fitness values that fitness functions map to is bounded by a polynomial then the corresponding fitness landscapes are easy.*

*Proof.* Any adaptive step increases the fitness value, and by transitivity, a fitness value is never repeated in an adaptive path. Thus, all adaptive paths are at most polynomially long: more specifically, bounded by one less than the number of distinct fitness values in the range of the fitness function.

Note that the above theorem does not put any constraints on what those fitness values are and thus depends only on the rank ordering (i.e. on the existence of very large rank equivalence classes). For variants that looks at the exact numeric values, see SA D.2 and Proposition 32.

The most notable application of Proposition 30 might be to the original Gavrilets and Gravner [6] formulation of holey adaptive landscapes. Gavrilets and Gravner [6] decided to divide the range of fitness values into just two components: 1 for viable and 0 for non-viable. An adaptive path in such a landscape can be at most 1 step long: if you start at a non-viable point right next to a viable one. Thus, all viability landscapes (like the original holey ones) are easy. Note that this does not mean fitness plateaus or neutral networks are easy to reach. If we remove the modeling restriction of just 0 or 1 fitness values and allow a large range of values then that does not eliminate the possibility of large fitness plateaus. In other words, fitness landscapes could have large plateaus of fitness but still be hard: i.e. it will be difficult to reach the plateau. Of course, more work should be done in the future on the topic of hardness of large neutral networks.

#### D.2 Approximate peaks

If we are to discuss neutral plateaus or networks then it is also important to consider nearly-neutral networks. For this, we need to use the whole numeric structure of the fitness function *f* and not just the rank-ordering that was sufficient until this point. Thus, let us consider relaxations of equilibrium, and being “close” to a peak instead of exactly at one. The following definitions and proofs are based on combinatorial optimization results by Orlin, Punnen, and Schulz [31].

**Definition 31.** A genotype *x* is at an *s*-approximate peak if *∀y ∈ N* (*x*) *f* (*y*) *≢* (1 + *s*)*f* (*x*).

The question becomes how big does *s* have to be for evolution to find an *s*-approximate peak. But since there is no absolute units of fitness, we will need to define *f*_*δ*_ = min_*x*_ min_*y*∈*N*(*x*) s.t. *f*(*y*)>*f*(*x*)_(*f* (*y*) *-f* (*x*)) and *f*_max_ = max_*x*_ *f* (*x*).

First, it is important to note that all landscapes where *f*_*δ*_ isn’t small compared to *f*_max_ are easy.

**Proposition 32.** *If f*_*max*_*/f*_*δ*_ ∈ *O*(*n*^*k*^) *for some constant k then an exact peak can be found in a polynomial in n number of mutations by any adaptive dynamic.*

*Proof.* Since each adaptive step increases fitness by at least *f*_*δ*_ then after *t* adaptive steps, we have *f* (*x*_*t*_) ≥ *f*_*δ*_*t*. Combine this with *f* (*x*_*t*_) ≤ *f*_max_ to get that *t* ≤ *f*_max_*/f*_*δ*_. □

So, we need to focus on bigger gaps between *f*_*δ*_ and *f*_max_. If the gap is exponential then we can find approximate peak for moderate sized *s* on any landscape.

**Theorem 33.** *If* log(*f*_*max*_*/f*_*δ*_) ∈ *O*(*n*^*k*^) *then fittest mutant SSWM dynamics can find a local s-approximate peak in time polynomial in n and* 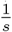.

*Proof.* Let *x*_0_ be the initial genotype, if it is an exact peak then we are done. Otherwise, let *x*_1_ be the next adaptive step, by definition of *f*_*δ*_, we have that *f* (*x*_1_) ≥ *f* (*x*_0_) + *f*_*δ*_ ≥ *f*_*δ*_. Now, consider an adaptive path *x*_1_*…x*_*t*_ that hasn’t encountered an *s*-approximate peak; i.e. a mutation was always available such that *f* (*x*_*i*+1_) > (1 + *s*)*f* (*x*_*i*_). Thus, we have that *f* (*x*_*t*_) ≤ *f*_max_ and that *f* (*x*_*t*_) ≥ (1 + *s*)^*t*^ *f*_1_ ≥ (1 + *s*)^*t*^ *f*_*δ*_. Putting these two together:

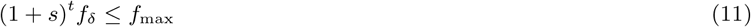

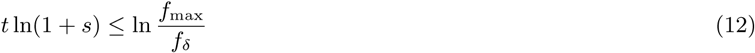

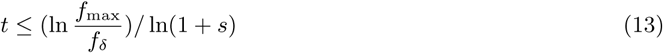

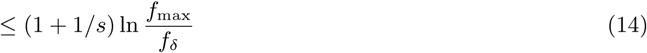

Where I used 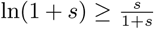 in the last step. Combining with the conditions on log *f_max_/f*_*δ*_, we get: 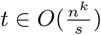. □

But for very small *s*, finding an approximate peak is as hard as finding an exact peak.

**Proposition 34.** *If s* ≤ *f*_*δ*_*/f*_*max*_ *then any s-approximate peak is a (exact) local peak.*

*Proof.* If an *s*-approximate peak at *x* is not an exact peak then there exists a *y* ∈ *N* (*x*) such that *f* (*y*) *-f* (*x*) ≥ *f*_*δ*_ but *f* (*y*) < (1 + *s*)*f* (*x*). Combining this with *f* (*x*) ≤ *f*_max_, we get that *s* > *f*_*δ*_*/f*_max_. □

Thus, it isn’t possible to find an *s*-approximate peak for very small *s* on hard fitness landscapes:

**Theorem 35.** *If PLS* ≠ *P and* log(*f*_*max*_*/f*_*δ*_) ∈ *O*(*n*^*k*^) *then (for NK-model with K* ≥ 2*) a local s-approximate peak cannot be found in time polynomial in n and* log 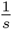.

*Proof.* If such an algorithm existed then we’d run it with *s* = *f*_*δ*_*/f*_max_ and – by Proposition 34 – the approximate peak it finds would be exact. Further, in this case log 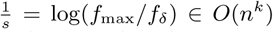 and thus the runtime would be polynomial in *n*. This is not possible for the NK-model with *K* ≥ 2 by Theorem 27 (unless PLS = P). □

This also means that the selective coefficient of the fittest mutant 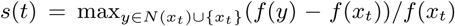 cannot decay exponentially quickly.

**Corollary 36.** *If PLS* ≠ *P then there are no evolutionary dynamics such that s*(*t*) ≤ *e*^*-mt*^ *for all instances of the NK-model with K* ≥ 2.

Contrast this with the always achievable power-law decrease in *s*(*t*).

#### D.3 Distributions and random fitness landscapes

The *NK* model is frequently studied through simulation, or statistical mechanics approaches. In a typical biological treatment, the gene-interaction network is assumed to be something simple like a generalized cycle (where *x*_*i*_ is linked to *x*_*i*+1_, *…x*_*i*+*K*_) or a random *K*-regular graph. The fitness contributions *f*_*i*_ are usually sampled from some choice of distribution. As such, we can think of biologists as doing average case analysis of these fitness landscapes. Given that randomly sampling landscapes can introduce structure like short paths [55], I suspect that the structure of this simple sampling led prior research to miss the possibility of exponentially long walks. The indepdent sampling of fitness components from the same distributions is especially apt to realize the conditions of Proposition 32 since it makes it unlikely to create an exponential gap between the smallest positive fitness gap *f*_*δ*_ and the maximum achievable fitness *f*_max_. Future work could provide a more careful analysis of this conjecture.

Given a historical disconnect between theory and data [25,53], the choice for distributions was usually made out of analytic convenience or (occasionally) out of the belief that a uniform distribution is akin to no assumption Since there is no strong empirical or theoretically sound justification for the choice of particular distributions of large fitness landscapes, I avoid relying on a simple generating distribution and instead reason from only the logical description of the model. This can be thought of as worst-case analysis or as analysis for arbitrary distributions of landscapes. By following this approach, we know that our results are features of the logic that characterizes a particular family of fitness landscapes and not artifacts of a particular simple sampling distribution. This is a standard method in theoretical computer science, but it is not as common in statistical physics or biology.

Although there is evidence for simple distributions on small fitness landscapes (on utpo 8 genes; see Szendro, Schenk, Franke, Krug, and de Visser [24] and Franke, Klozer, de Visser, and Krug [60]), there is little to no data on the distribution of large (i.e. on many loci) fitness landscapes in nature. And as discussed in the main text, given the exponential size of fitness landscapes, it is unlikely that such data could be collected. However, if a single sampling distribution is required then it is tempting to turn to Occam’s razor and consider simpler landscapes as more likely. This can be done by sampling landscapes with negative log probability proportional to their minimum description length, i.e. according to the Kolmogorov universal distribution. If landscapes are sampled in this way then I would expect all the orders of magnitude for hardness results established herein to hold [56]. However, I leave it as an open question for future work to prove this formally and to contrast the ubiquity of hardness in fitness landscapes sampled under different theoretical distributions. As outlined above, it would be especially interesting to analyze the uniform distribution (that is popular in statistical physics) versus the Kolmogorov universal distribution (that is used in theoretical computer science).

